# Dynamic Workforce Modulation and Foraging Efficiency in Eusocial Insect Colonies

**DOI:** 10.64898/2026.04.21.719794

**Authors:** Daniel Campos, Javier Cristín, Pol Fernández-López, Frederic Bartumeus

## Abstract

Understanding the fitness advantages conferred by eusociality remains a central challenge in behavioral ecology. One promising approach is to identify collective strategies that shift efficiency within social groups. Here, we test the hypothesis that reserve workforces in eusocial insect colonies represent an adaptive mechanism that enhances flexibility and foraging efficiency under fluctuating environmental conditions. We examine how such reserve workers modulate the departure and return rates of foragers and how these time-dependent dynamics shape the colony’s overall energetic balance. By integrating an energetic-balance framework with stochastic search simulations inspired by empirical results from *Aphaenogaster senilis*, we quantify the energetic requirements for colony viability, incorporating energy intake, search costs, and basal metabolic demands. Our results show that as colonies grow, maintaining a positive energy balance requires a disproportionately larger relative workforce. By modulating departure and return rates over time, colonies control the synchrony of their collective search and efficiently activate or suppress their reserve workforce to scale foraging effort as needed. These findings suggest that the “lazy” or weakly engaged workers commonly observed in large colonies function as an essential reserve that stabilizes colony energetics and enhances responsiveness. Together, our results provide a functional explanation for sublinear metabolic scaling in eusocial groups and highlight workforce modulation as a key factor underlying their energetic stability and evolutionary success.

## I. INTRODUCTION

Eusociality provides biological species with a fundamentally expanded repertoire of adaptive strategies, offering substantial advantages in ecological and evolutionary contexts. This view is reinforced by genetic evidence showing that such a complex form of social organization has arisen independently and repeatedly throughout evolutionary history (see [1–6] for recent accounts). Additionally, phylogenetic methods provide evidence that the transition from solitary to collective strategies, resulting from colony growth/development is rarely reversed [4]. As a central component of this expanded adaptive repertoire, dynamic task allocation and colony-level activation have been experimentally shown to rely on regulatory mechanisms that respond to external fluctuations or disturbances, thereby providing colonies with substantial behavioral flexibility [7]. In line with this, ant colonies can shift from individual to collective foraging strategies depending on resource size [8], density [9–12], and quality [13]. Moreover, increases in group size typically favor collective over individual foraging strategies [13, 14].

However, eusociality also entails that the intrinsic value of any given individual decreases as group size increases. Intriguingly, this promotes a form of redundancy within colonies, whereby active workers coexist with inactive individuals that remain available for rapid recruitment when specific tasks arise [15]. The proportion of these “inactive” or “lazy” workers has been shown to increase with colony size [16–21]. Although maintaining such inactive individuals entails non-trivial metabolic costs, several hypotheses have been proposed to explain their persistence [20]. The most strongly supported view interprets these workers as a reserve workforce that equips the colony with the capacity for rapid adaptive responses to transient demands or extreme events [22, 23].

More broadly, the use of redundant search units—formalized as the redundancy principle [24]—has been proposed as an evolutionary adaptation across a wide range of biological and chemical systems [25–31]. The central idea is straightforward: increasing the number of searchers, whether molecules or organisms, reduces the expected search time for targets, and natural selection should therefore favor redundancy whenever feasible. However, in the context of social foraging, energetic constraints become critical, as deploying an arbitrarily large workforce may be metabolically inefficient. This trade-off suggests that colonies must regulate how many workers are active at any given time, making the dynamic modulation of task-related workforces a potentially essential component of the transition from individual to collective behavior. It also highlights the importance of understanding how colonies balance the benefits of redundancy with the energetic costs of maintaining and activating additional workers. Although empirical evidence is still limited, several studies indicate that such modulation enables colonies to maintain—or rapidly restore—overall activity levels even after the removal of a substantial fraction of active workers. This compensatory response has been documented in Temnothorax rugatulus [16], Myrmica kotokui [32], and in bees [33]. Quantifying the specific benefits that workforce modulation and the emergence of redundancy provide to ant colonies therefore represents a key step toward understanding the functional basis of eusociality.

In the present work, we (i) provide numerical and empirical evidence that dynamic workflow allocation acts as a convenient—and potentially necessary—mechanism conferring additional behavioral flexibility to eusocial groups, and (ii) examine the conditions under which such enhanced flexibility compensates for the metabolic costs associated with maintaining inactive workers. More specifically, we investigate how the time-dependent modulation of reserve workforces alters colony-level exploration patterns—such as the number of simultaneous searchers and the average distances travelled by active foragers—and how these changes affect search efficiency, quantified through the energetic balance resulting from foraging activity.

Mathematically, previous work has show that for a group of *N* diffusive individuals starting their search at *t* = 0 from a common place (e.g. the nest), the time required to reach a single target located at distance *d*_*r*_ from the nest scales linearly with 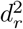 and decays logarithmically with *N*, provided these variables are large enough (see [34–39] for a survey). Here, we extend this classical framework by considering a more realistic scenario in which the number of active searchers varies in time. We denote by *N*_*a*_(*t*) the number of active workers, which become active or inactive at rates *γ*_*b*_ and *γ*_*d*_, respectively (with subscripts *b* and *d* referring to “birth” and “death” of search trajectories). In this formulation, the limit *γ*_*b*_ →0 corresponds to highly asynchronous search, where individuals leave the nest sporadically, whereas *γ*_*b*_ →∞ represents fully synchronous collective search. Once a searcher completes a trajectory (at rate *γ*_*d*_) it becomes inactive again (effectively returning to the nest) so individuals continuously switch between active and inactive states until the search concludes. In previous work [40], we identified the optimal values of *γ*_*b*_ and *γ*_*d*_ that minimize the collective search time required to reach the target. In the present study, we substantially extend this idea by incorporating energetic gains and costs relevant to eusocial colonies. Our goal is to determine under which conditions the presence of reserve workforces becomes particularly advantageous, and how the interplay between activation rates, deactivation rates, and energetic constraints shapes the efficiency of collective search.

## II. METHODS

### A. Empirical data acquisition and processing

In order to explore how ant colonies arrange and regulate workforces during a search, we used experimental data obtained in the lab (under constant temperature conditions at 20ºC) from two different experiments, each conducted with two different colonies of *Aphaenogaster senilis* ants.

#### Experiment A

First, we conducted an experiment in an arena consisting of a set of consecutive parallel (leftwards and rightwards) lines, so reproducing a quasi-1d topological space (see Fig. 1a). The nest was connected through a plastic tube to one end of the structure, and the lines were surrounded by water to prevent the ants to escape, or get out from the structure. Food (consisting of small amounts of cookies and egg yolk) was located 1 meter away from the nest entrance. Then, three different food conditionings were considered by either adding new food items at 2 meters from the nest (high-density condition), at 3 meters away from the nest (medium-density condition), or non adding any additional resources (low-density condition).

**FIG. 1.**
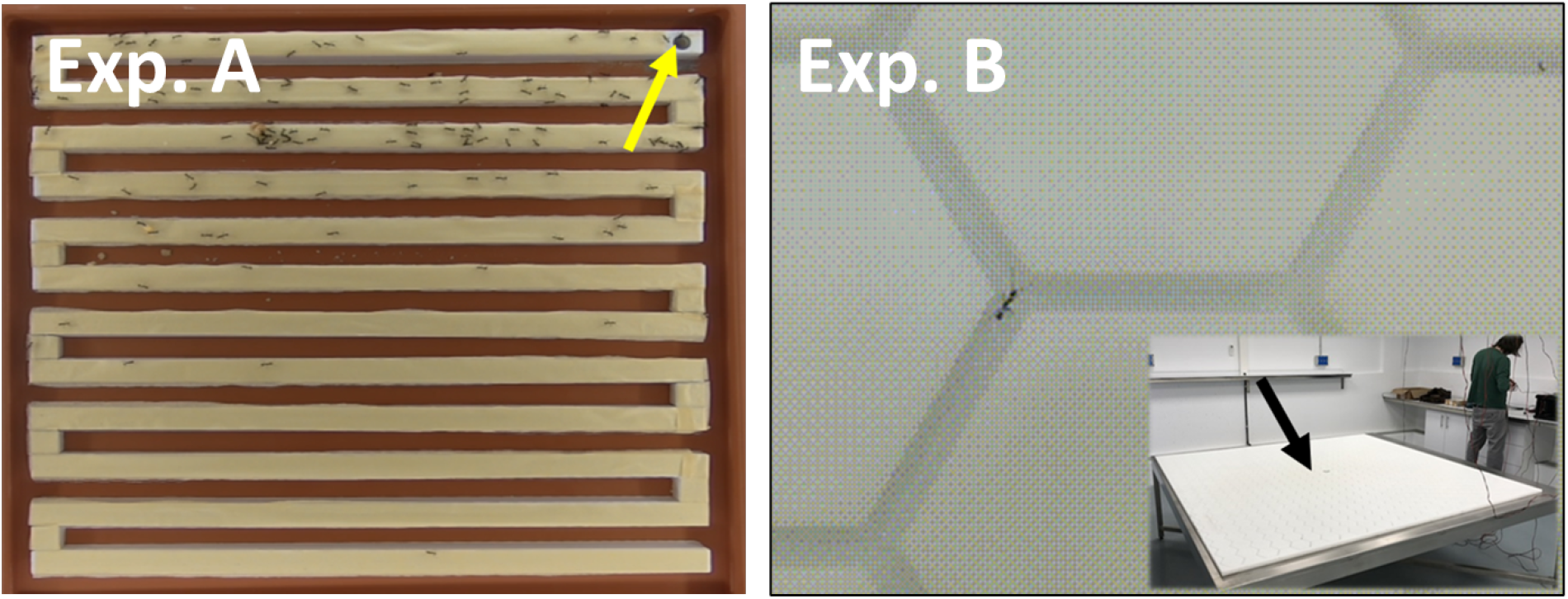
Illustration of the different arenas used to obtain the experimental data used in the work (the arrows identify the entrance to the nest in each case). Experiment A: Quasi-1d arena consisting of parallel lines leftwards and rightwards. Experiment B: Hexagonal lattice (the main image shows a close-up of the lattice, while the inset on the lower right corner corresponds to the whole structure).

In this experiment we compared the foraging dynamics of two colonies of different size (designated colonies A1 and A2 in the following). Colony A1 had *N*_*col*_ = 200 individuals and colony A2 *N*_*col*_ = 60 approximately (these numbers did not vary significantly during the experiments). Three experiments per week (in nonconsecutive days) were conducted for each colony until completing a total of five experiments for each colony and each food conditioning.

#### Experiment B

Additional experimental datasets were obtained from a previous setup [41] corresponding to two other colonies of approximately the same size (*N*_*col*_ ≈750 individuals), named B1 and B2. These were allowed to explore a large arena consisting of a hexagonal lattice with a total size of 2x1 meters (see Fig 1b). The details about this second setup and dataset have already been given and discussed in [41]. Three food conditions were considered for this case: (i) deterministic, (ii) stochastic, and (iii) no-food. In the deterministic case food resources were located at two nodes (always the same) in the lattice at a medium distance from the nest, while in the stochastic case the recources were located at two different nodes selected at random every day.

In both experiments A and B the foraging process (starting when the arena is connected to the nest) was recorded in video for a period of 2 hours, which was far enough for the ants to find and collect the food, and to explore the rest of the arena. Actually, the period of food gathering typically elapsed about 30-50 minutes. The recording was done at a frequency of 10 Hz, and the analysis of the frames was carried out with standard libraries (in Matlab). As a result, we could compute the number of active ants present in the arena during the whole foraging process, *N*_*a*_(*t*).

From the data for *N*_*a*_(*t*) we could estimate average values of the rates *γ*_*b*_ and *γ*_*d*_. For this, we considered a moving time window of size *τ* throughout the experiments, and we computed the number of search trajectories appearing and terminating within the window (*n*_*in*_ and *n*_*out*_, respectively). According to the definition above, the average rate at which search trajectories appear is given by *γ*_*b*_ = *n*_*in*_*/τ* . Similarly, for all the ongoing trajectories present in the arena we have that they can be terminated at a rate *γ*_*d*_ according to our definition, so we use *γ*_*d*_ = *n*_*out*_*/* ⟨*N*_*a*_⟩ _*τ*_ *τ* (where ⟨*N*_*a*_⟩ _*τ*_ represents the number of active ants in the arena averaged over the time window *τ*). At practice, we took *τ* = 10 min, which was large enough to obtain significant statistics during the time window, while still low enough to let us observe an evolution of the foraging strategies through time.

### B. Computational model and parameters

We used a stochastic search model where searchers carry out random walks on an infinite plane starting from a common point (representing the nest). At time *t* = 0 a first search trajectory starts from the nest, and subsequently new random-walk trajectories appear (disappear) at rates *γ*_*b*_ (*γ*_*d*_), respectively. The corresponding search process is kept until the resources, located at a single random patch at a distance *d*_*r*_ from the nest, are hit by a trajectory (this happens if the distance from the trajectory to the resources is lower than a detection distance *d*_*d*_).

For simplicity, we assume that random walks obey classical Brownian motion, so each trajectory consists of a combination of independent Gaussian jumps with a characteristic speed *v* in random uncorrelated directions. Then, the length covered in a single jump during a time step Δ*t* is *v*Δ*t*. Accordingly, Δ*t* represents the characteristic persistence time of the trajectories, so for times shorter than Δ*t* the motion is persistent, while for times much longer than Δ*t* the direction of motion becomes uncorrelated. We will focus in the low-persistence regime given by Δ*t* ≪ *d*_*r*_*/v*, which at practice means that a large number of jumps is required to cover the distance *d*_*r*_ from the nest to the target, and so diffusive scaling will dominate at the relevant timescales for the search process.

In summary, the algorithm of the model used takes the form

For each time step *t* → *t* + Δ*t*, do:

i. *With probability γ*_*b*_Δ*t a new trajectory is initiated at the nest position*.
ii. *For every present foraging trajectory:*

iia) *With probability γ*_*d*_Δ*t, terminate the search trajectory*.

iib) *With probability* 1 −*γ*_*d*_Δ*t, add a new Gaussian displacement (with standard deviation v/*Δ*t) to the present position of the trajectory*.

iic) *If the resulting position is within a distance d*_*d*_ *to the resource location, then terminate the search*.

Note that the number of simultaneous search trajectories in the system will change with time. If we denote this number by *N*_*a*_(*t*), our initial condition states that *N*_*a*_(0) = 1, and then *N*_*a*_(*t*) will either increase or decrease until reaching the stationary value *γ*_*b*_*/γ*_*d*_ on average. At practice, for *γ*_*b*_ ≫ *γ*_*d*_ the number of searchers will increase to reach *N*_*a*_(*t*) = *γ*_*b*_*/γ*_*d*_ and then it will fluctuate around this value. Then, by denoting the number of ants in the colony as *N*_*col*_, and assuming that only a part *N*_*w*_ *< N*_*col*_ of them is susceptible to become recruited for foraging, the condition *γ*_*b*_*/γ*_*d*_ ≤ *N*_*w*_ must be satisfied, which sets a limitation on the range of *γ*_*b*_, *γ*_*d*_ values that are accessible to the colony. For the contrary case, *γ*_*b*_ ≪ *γ*_*d*_, most of the time we will have either just one single search trajectory (*N*_*a*_(*t*) = 1) or the search space will be empty (*N*_*a*_(*t*) = 0), and so the colony will rely on an *individual* foraging strategy.

As an output of the model, we computed the Search Time *T*_*fp*_ (this is, a first passage time, the time it takes for at least one of the trajectories to hit the resources for the first time), and the Collective Time Cost *T*_*c*_, defined as the combined duration of all trajectories that have emerged from the nest up to time *T*_*fp*_.

### C. Estimation of the energy balance in the colony

Since the main objective of the model above is to estimate whether an increasing workforce *N*_*w*_ available is beneficial or not for the colony, we need to estimate the energy intake and costs related to the search. There is a large controversy on what is the biologically relevant magnitude (e.g. the global energy intake, the energy per unit time, etc) that organisms or groups should optimize to maximize its fitness [42]. For the context considered in our work, we will assume that colonies should try to reach a positive balance between their energy consumption (which includes the basal consumption and the energy cost of foraging) and the energy intake obtained through foraging.

In the following, we describe how the energy balance can be conveniently written in terms of the parameters *N*_*col*_, *N*_*w*_, *T*_*fp*_ and *T*_*c*_ introduced above. For this, we assume that ants in the colony have always a constant basal metabolic rate RM_*b*_, and active foragers have an additional working metabolic rate RM_*a*_, so active foragers have a total metabolic rate RM_*b*_ + RM_*a*_. Also, we introduce *T*_*f*_ as the daily time that the colony spends foraging (so *T*_*f*_ *< T*_*day*_, with *T*_*day*_ representing the total duration of one day). Given our definition for the time required to find a food item, *T*_*fp*_, the average number of items that can be obtained by the colony in a single day is then *T*_*f*_ */* ⟨*T*_*fp*_⟩ (note that we use the bracket notation ⟨·⟩ to denote averages over the random trajectories, and similarly we will use ⟨·⟩_*t*_ to denote time averages).

Taking into account all these considerations, the different terms of the energy balance read:

1. **Energy intake**. If we denote the energy content of every food item as *E*_*item*_, the daily intake obtained by the colony as a result of the foraging process is given by

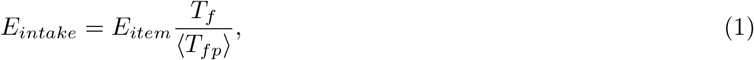

where we are not explicitly considering the gathering time (that is, the time it takes to bring the food items to the nest), but only the time required to find the food.
2. **Basal consumption**. According to the definitions above, the basal consumption for the whole colony of size *N*_*col*_ will be denoted by

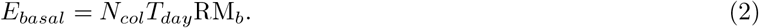
3. **Foraging cost**. The foraging cost will take into account the average number of active workers, *N*_*a*_, during the foraging time *T*_*f*_ . So that, it can be written as

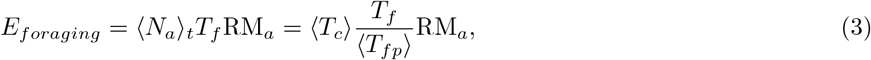

according to the definition of *T*_*c*_ given above, from which ⟨*T*_*c*_⟩ = ⟨*N*_*a*_*T*_*fp*_⟩_*t*_ ≈ ⟨*N*_*a*_⟩_*t*_⟨*T*_*fp*_⟩ follows.

In consequence, the energy balance for the colony takes the form

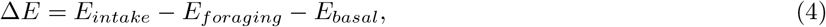

which can be conveniently rewritten as

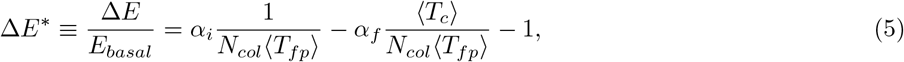

after introducing the parameters *α*_*i*_ ≡ *E*_*item*_*T*_*f*_ */T*_*day*_RM_*b*_ and *α*_*f*_ ≡ *T*_*f*_ RM_*a*_*/T*_*day*_RM_*b*_. These parameters *α*_*i*_ and *α*_*f*_ then represent the relative cost of *intake* and *foraging*, respectively. Since the metabolic rates and the rest of parameters involved in their definition are case and specie specific, and often they are not available experimentally, here we will rather explore the behavior of Δ*E*^*^ (by running random-walk simulations to compute ⟨*T*_*fp*_⟩ and ⟨*T*_*c*_⟩) as a function of *α*_*i*_ and *α*_*f*_, with the focus put on identifying the region where the balance Δ*E*^*^ becomes positive, which determines the condition for the colony to become energetically viable.

In particular, the questions we will address in the present work are: given some values of *α*_*i*_ and *α*_*f*_, what are the combinations of *γ*_*b*_, *γ*_*d*_, *N*_*col*_, *N*_*w*_ values (with the aforementioned constraint *γ*_*b*_*/γ*_*d*_ ≤ *N*_*w*_) that lead to Δ*E*^*^ *>* 0? When is it convenient for the colony to increase its workforce *N*_*w*_ as a mechanism to increase its energy intake, even at the expense of increasing its foraging cost?

## III. RESULTS AND DISCUSSION

### A. Time evolution of active foragers in the arena

We quantified the time evolution of departures (birth rate, *γ*_*b*_) and returns (death rate, *γ*_*d*_) of foraging ants in the two experimental arenas (Fig. 2). Colonies exhibited a gradual increase in the number of ants entering the arena, a pattern driven by rising birth rates—more or less pronounced depending on the colony and the experiment—and by consistently decreasing death rates. The latter aligns with a strategy commonly reported in many ant species: exploration begins with one or a few scouts that assess external conditions and help the colony evaluate potential risks outside the nest. Only once safety is established does the number of searching individuals, and the diversity of their trajectories, expand. Our goal in this work is to determine whether, and how, these temporal trends in activation and deactivation influence the colony’s overall energetic balance.

**FIG. 2.**
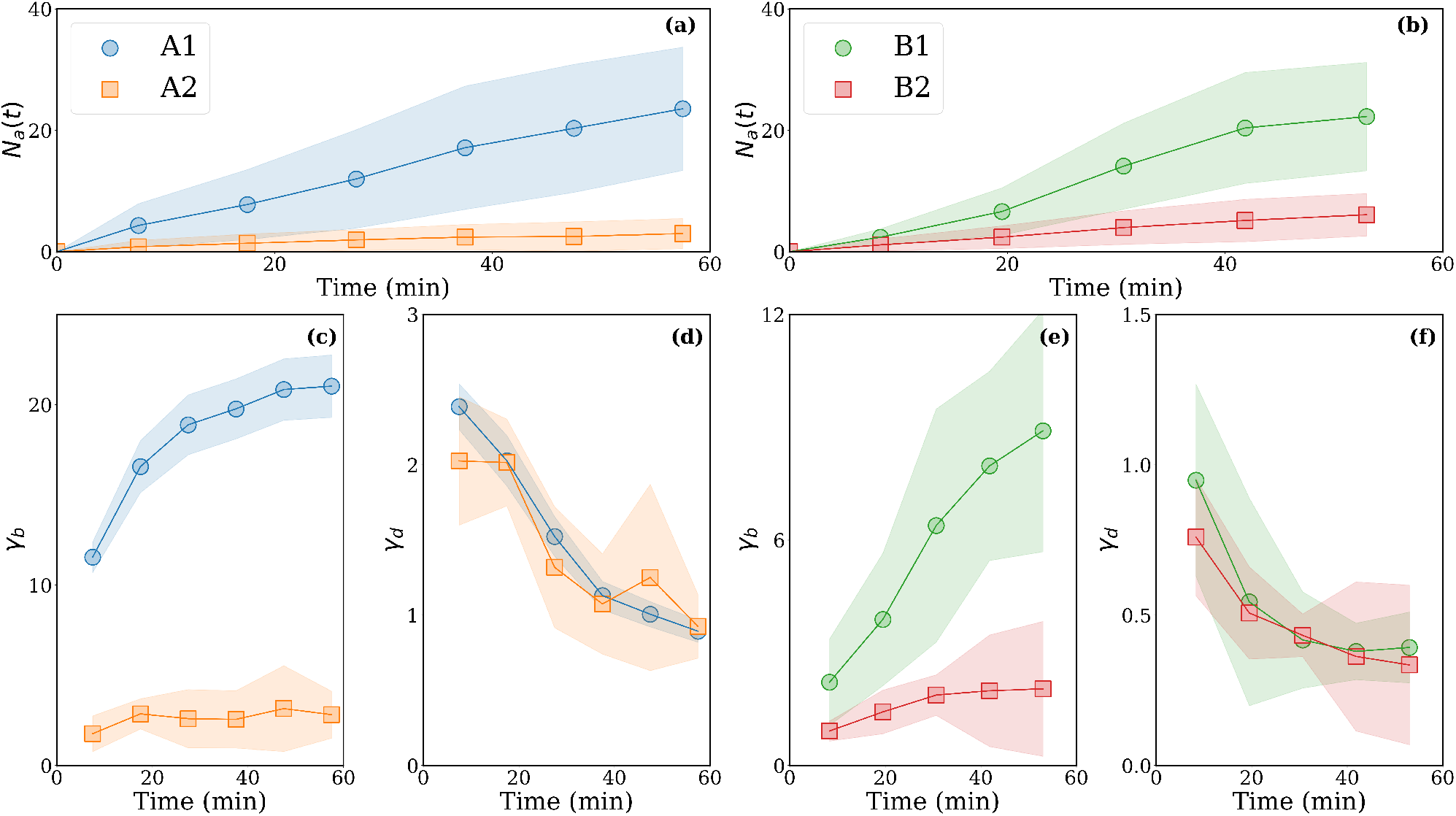
Average number of ants *N*_*a*_(*t*) and rates *γ*_*b*_ and *γ*_*d*_ obtained from the experimental datasets for the four colonies considered, two for experiment A (left) and two for experiment B (right). Only data from the first 60 minutes of the foraging process is considered, which corresponds in most cases to the search phase and the initial part of the food recollection phase. Error bars (shaded region) correspond to the standard deviation obtained by averaging over all the daily experiments carried out for each colony.

Experiment A (Fig. 2a,c,d) allows us to compare the foraging strategies (identified through the dynamics of rates *γ*_*d*_ and *γ*_*b*_) used by two relatively small colonies of different size. It is seen that there is a clear tendency of *γ*_*b*_ (*γ*_*d*_) to increase (decrease) throughout the foraging process. Furthermore, the larger colony (A1) also exhibits a higher activity in terms of the number of active workers (note that *N*_*a*_ for colony A1 is about six times that for colony A2, though its size *N*_*col*_ is only ≈2.5 times larger). Finally, it is also evident that the larger activity of colony A1 is orchestrated through a much larger number of search trajectories (evidenced through larger birth rates *γ*_*b*_), while durations of these trajectories are approximately the same for both colonies (this is, their *γ*_*d*_ rates do not vary significantly).

Colonies B1 and B2 in experiment B (Fig. 2b,e,f), have approximately the same size. However, we observed a large variability between their foraging activities throughout the experiments (the number of active foragers *N*_*a*_ was on average two times larger for B1). This may be due to the history of the two colonies (they may be originally adapted to different conditions before the experiments, or their task partitioning/needs may be different). Whatever the specific reason, different foraging activities *N*_*a*_(*t*) may presumably reflect different workforces *N*_*w*_, too.

In experiments B, we found similar results for the rates despite using a different arena and food conditions. We identified a clear tendency to increase *γ*_*b*_ values and decrease *γ*_*d*_ values with time during the foraging process; this seems to be a rather robust dynamics through all colonies and conditions considered. Besides, the difference in foraging activities between colonies B1 and B2 results again from differences in *γ*_*b*_ (rather than *γ*_*d*_, which shows similar values for both colonies).

We note that although Fig. 2 only shows averaged rates, the intraday (morning *vs*. afternoon) or interday variability observed between experiments does not significantly affect our conclusions above (in Appendix A we provide and discuss the corresponding results for the sake of completeness). Extending our results to later phases of foraging (gathering and pos-gathering stages) shows that once the entire arena has been explored and no additional food resources are available, the colony tends to progressively decrease the number of active workers in the arena, reflected by a slight increase in *γ*_*d*_. These trends, which are also presented in Appendix A Figs. 6 and 7, are mostly evident in colonies B1 and B2, while colonies A1 and A2 did not reach such a decline in activity during the duration of the experiments.

Finally, we stress that food conditioning does not seem to induce any significant trend or variation in the strategies of the different colonies, at least in terms of *γ*_*b*_ and *γ*_*d*_. This is true even in the case of colonies B1 and B2, where food amount and disposition in the arena was kept the same for two/three consecutive weeks of experiments [41]. So, we find no evidence of adaptation to resource distributions throughout the experiments. This may be because the differences in food spatial allocation used in our experiments were not sufficiently large. An alternative explanation would be that the adaptation of collective strategies to resource distributions, if any, requires longer (maybe pluriannual) monitoring of the colony’s foraging patterns. Specific experiments should then be necessary to confirm or discard these ideas.

### B. Energy balance optimization of collective searches through *γ*_*b*_ and *γ*_*d*_

In Fig 3, we show heatmaps of the average energy balance Δ*E*^*^. The quantity Δ*E*^*^ is computed through random-walk search simulations that allow us to determine ⟨*T*_*fp*_⟩ and ⟨*T*_*C*_⟩ (see Appendix B) as functions of the departure and return rates *γ*_*b*_ and *γ*_*d*_, and for a specific choice of the parameters *α*_*i*_ and *α*_*f*_ . The values of the parameters *α* were selected to ensure that we explore the space parameter around Δ*E*^*^ ≈0, so that we have a clear transition between a positive and a negative energy balance. Alternative choices of *α*_*i*_ and *α*_*f*_ were also explored, and we verified that they do not qualitatively alter our results.

**FIG. 3.**
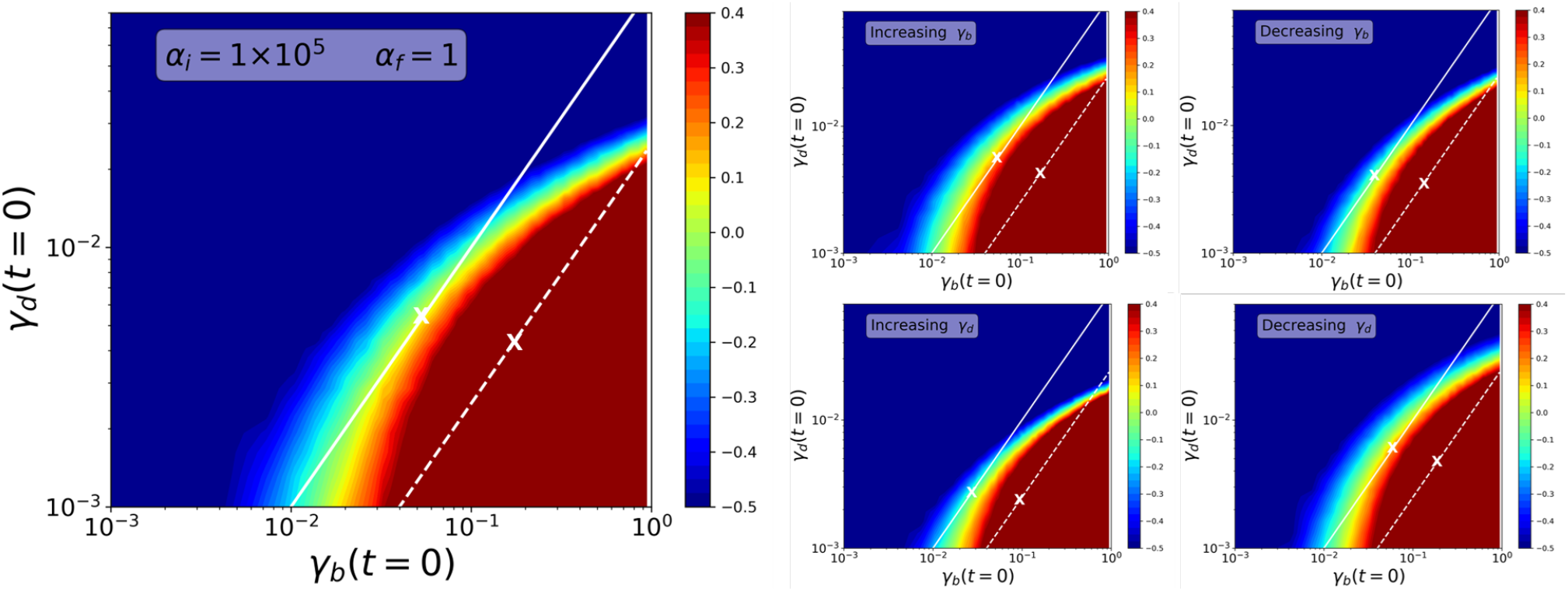
Heatmaps obtained from random-walk simulations for the energy balance Δ*E*^*^ as a function of rates *γ*_*b*_, *γ*_*d*_ (using *d*_*r*_ = 10, *D* = 1, *d*_*d*_ = 1). (Left): reference case for specific values of *α*_1_, *α*_2_ (indicated in the legend), chosen so the shape of the region Δ*E*^*^ *>* 0 where the colony would be energetically viable can be identified. (Right): case when *γ*_*b*_(*t*) or *γ*_*d*_(*t*) are allowed to increase or decrease by a factor of two during the search. Such an increase or decrease is linear with time and terminates after a characteristic time 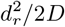, so after that time *γ*_*b*_ and *γ*_*d*_ will remain constant for the rest of the simulation. Solid and dashed lines correspond to the upper bounds *N*_*w*_ = 0.05*N*_*col*_ and *N*_*w*_ = 0.2*N*_*col*_, respectively, which mean that only regions on the left of such lines are accessible, provided that only 5% or 20% of the colony individuals can get engaged in foraging tasks. The maximum value of Δ*E*^*^ for such regions is indicated with an ‘x’ (note that for some cases this point lies in a region where Δ*E* does not change significantly, and so the heatmap shows a relatively constant colour).

Inspired by our experimental results (showing increasing *γ*_*b*_ and decreasing *γ*_*d*_ over time), we also analysed how time-dependent departure and return rates affected the energy-balance parameter space. Specifically, we allowed either *γ*_*b*_ or *γ*_*d*_ to increase (or decrease) from an initial value, denoted as *γ*_*b*_(*t* = 0) and *γ*_*d*_(*t* = 0), until reaching twice (or half) that value at time 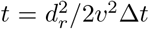, which is on the order of the expected search time required to find a food item located at distance *d*_*r*_. After this point, the rates were kept constant at the values attained. This procedure allowed us to explore how gradual adjustments in departure and returning rates, including those consistent with the temporal patterns observed experimentally, reshape the colony’s energetic balance landscape.

We observe that more favourable energetic situations—that is, higher values of (Δ*E*^*^), consistently cluster in the region of large *γ*_*b*_ and small *γ*_*d*_. Accordingly, Fig. 3 (right) shows that either increasing *γ*_*b*_ or decreasing *γ*_*d*_ leads to more favourable values of Δ*E*^*^ compared with the reference case (large panel on the left), where both rates remain constant. Notably, this pattern, increasing departures and decreasing returns with time, is precisely the trend shared by all colonies in the experimental data discussed in the previous section. Conversely, increasing *γ*_*d*_ or decreasing *γ*_*b*_ shifts the system toward less favourable energetic outcomes.

In Appendix B we show that, according to the energy balance in Eq. 5, large workforce loads and synchronous individual searches reduce first-passage times *T*_*fp*_ while keeping collective search costs relatively high. Nonetheless, in resource-limited scenarios, where resources are far away or difficult to detect, finding and collecting the food takes more time and the collective search cost *T*_*C*_ of many synchronous search trajectories becomes too high and might be reduced exploiting more asynchronous search strategies, where ant workers depart the nest at slower rates. The fact that, in our simulations, the most favourable energetic balances (Fig. 3) are obtained for large *γ*_*b*_ and small *γ*_*d*_ simply reflects that the search time (*T*_*fp*_) dominates the energetic balance over *T*_*C*_. In our search initial condition, engaging an increasingly large number of individuals in the search process would be the optimal strategy.

However, our simulations do not explicitly incorporate the costs or risks of predation, nor other detrimental effects that such long-time massive collective deployment may entail. We can indirectly account for these effects by assuming that the colony has access to a maximum workforce (*N*_*w*_). Given the condition (*γ*_*b*_*/γ*_*d*_ ≤ *N*_*w*_), this is equivalent to imposing an upper bound on the ratio *γ*_*b*_*/γ*_*d*_. In the heatmaps, we therefore indicate the region of *γ*_*b*_)*and*(*γ*_*d*_) values available to the colony under the assumption that only a fraction (*N*_*w*_*/N*_*col*_) of individuals can be engaged in foraging. These accessible regions correspond to the areas to the left of the solid and dashed lines for the cases (*N*_*w*_*/N*_*col*_ = 0.05) and (*N*_*w*_*/N*_*col*_ = 0.2), respectively.

The interpretation is straightforward: if, for example, only (5%) of the colony can be allocated to foraging—due to risks or because the remaining workers must perform other tasks—then only the region to the left of the solid line is attainable. Within this constraint, the colony would maximise its fitness by occupying a point in that region where Δ*E*^*^ *>* 0. For completeness, we mark in the heatmaps the specific positions where (Δ*E*^*^) reaches its maximum under the *N*_*w*_ restriction (denoted by “x” symbols). Notably, these optima typically occur at nontrivial intermediate values of *γ*_*b*_ and *γ*_*d*_.

### C. Energetic balance as a function of the relative workforce and colony size

Next, we analyze in more depth whether the availability of the workforce *N*_*w*_ constrains the potential foraging activity of a colony and thus influences its energetic balance. First, in Fig. (4) we show the maximum energy balance 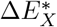 (this is, the energy balance reached at the ‘x’ points of Fig. 3) as a function of the foraging workforce available (*N*_*w*_) for colonies of different sizes (*N*_*col*_ = 100 and *N*_*col*_ = 500, corresponding to small and large circles in the plots, respectively). We compare the reference case (*γ*_*b*_ and *γ*_*d*_ constant, empty circles) to the more favourable case corresponding to ‘increasing *γ*_*b*_’ (full circles), where the latter, as expected, reaches always higher 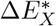 values.

**FIG. 4.**
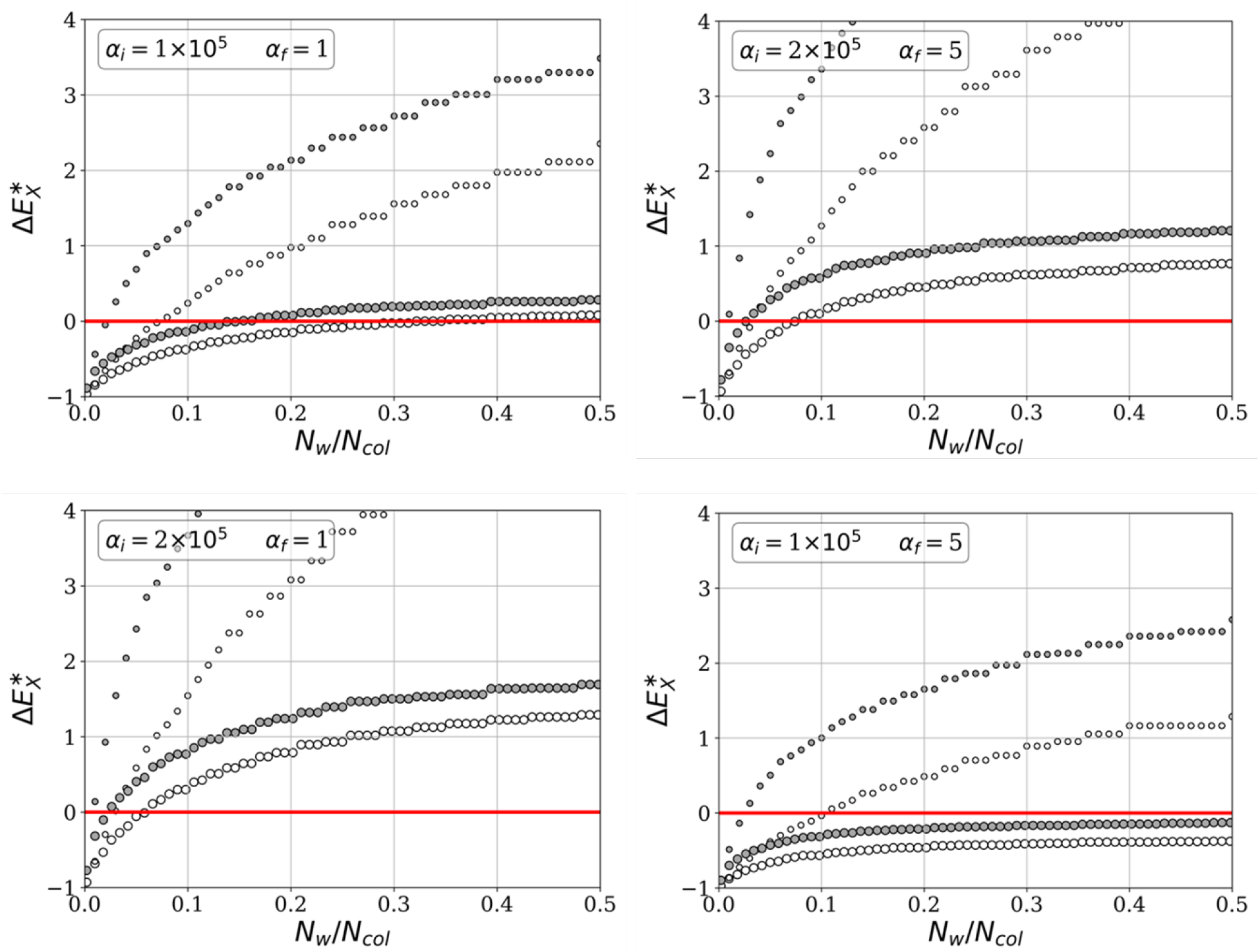
Maximum energy balance that can be reached by a colony of size *N*_*col*_ = 100 (small circles) and *N*_*col*_ = 500 (large circles) as a function of the foraging workforce *N*_*w*_ (expressed as a fraction of *N*_*col*_). Empty circles represent the cases with *γ*_*b*_ and *γ*_*d*_ constant, and black circles show the case with increasing *γ*_*b*_. Different values of *α*_1_ and *α*_2_ are considered (indicated in the plot legends).

Fig. (4) shows consistently (i.e., for different choices of *α*_*i*_ and *α*_*f*_ values) that for a small colony under the same search conditions, foraging becomes much more cost-effective. The viability of the large colony is always compromised in comparison to that of the small colony, as 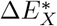 values for the former are always lower. Actually, we identify situations (see case *α*_*i*_ = 10^5^, *α*_*f*_ = 5) where the large colony is not viable (i.e., 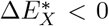 for any workforce *N*_*w*_), while the small colony is.

Even in cases where both colonies can achieve a positive energetic balance, there is always a minimum workforce *N*_*w*_ required for such energetic viability to be reached. This highlights that having a sufficiently large workforce represents a significant constraint for these colonies. Moreover, this requirement becomes increasingly restrictive as colony size grows. This can be seen by noting that the minimum workforce needed to obtain a positive energy balance is smaller for the case *N*_*col*_ = 100. In conclusion, small colonies appear to be less limited by the amount of workforce required to collect food in an energetically efficient manner.

To complete our understanding of how energetic balances constrain ant colony foraging processes, Fig. 5, shows how the optimal values of *γ*_*b*_, *γ*_*d*_ (those reached at the ‘x’ points, denoted by (*γ*_*b*_)_*X*_ and (*γ*_*d*_)_*X*_) vary as functions of the colony size and the available workforce.

**FIG. 5.**
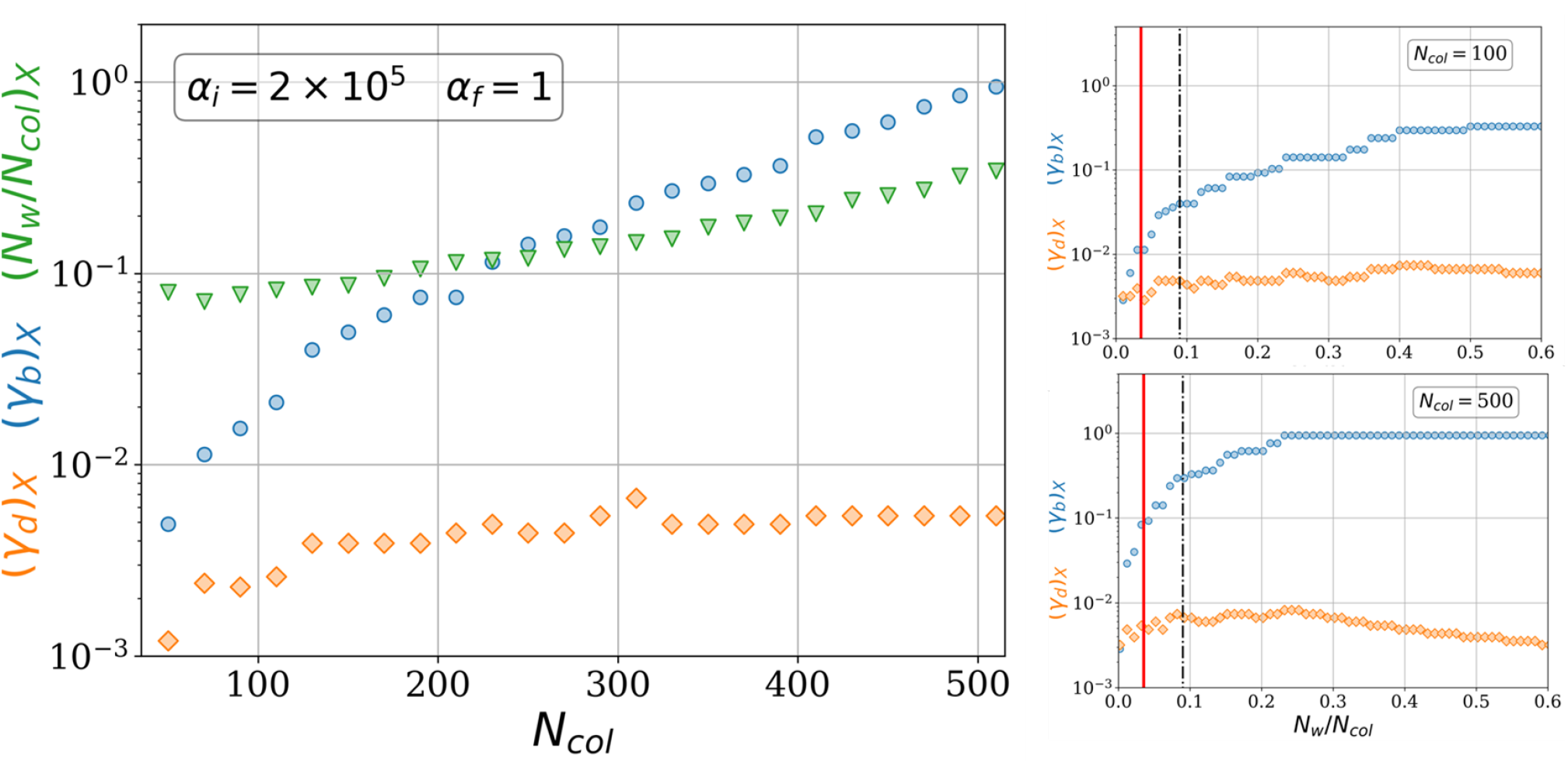
Left: Rates *γ*_*b*_ (blue circles) and *γ*_*d*_ (orange diamonds) corresponding to the maximum energy balance 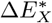 as a function of the colony size. The relative workforce *N*_*w*_*/N*_*col*_ corresponding to that optimal balance in every case is also shown (green triangles). Right: The same rates *γ*_*b*_ and *γ*_*d*_ are plotted for two colony sizes *N*_*col*_ = 100 and *N*_*col*_ = 500 (see legend) as a function of the foraging workforce *N*_*w*_ (expressed as a fraction of *N*_*col*_). The vertical red solid line identifies the minimum workforce corresponding to 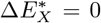, while the black dashdot line identifies the optimal workforce value reached when the energy balance reaches a maximum 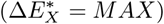.

**FIG. 6.**
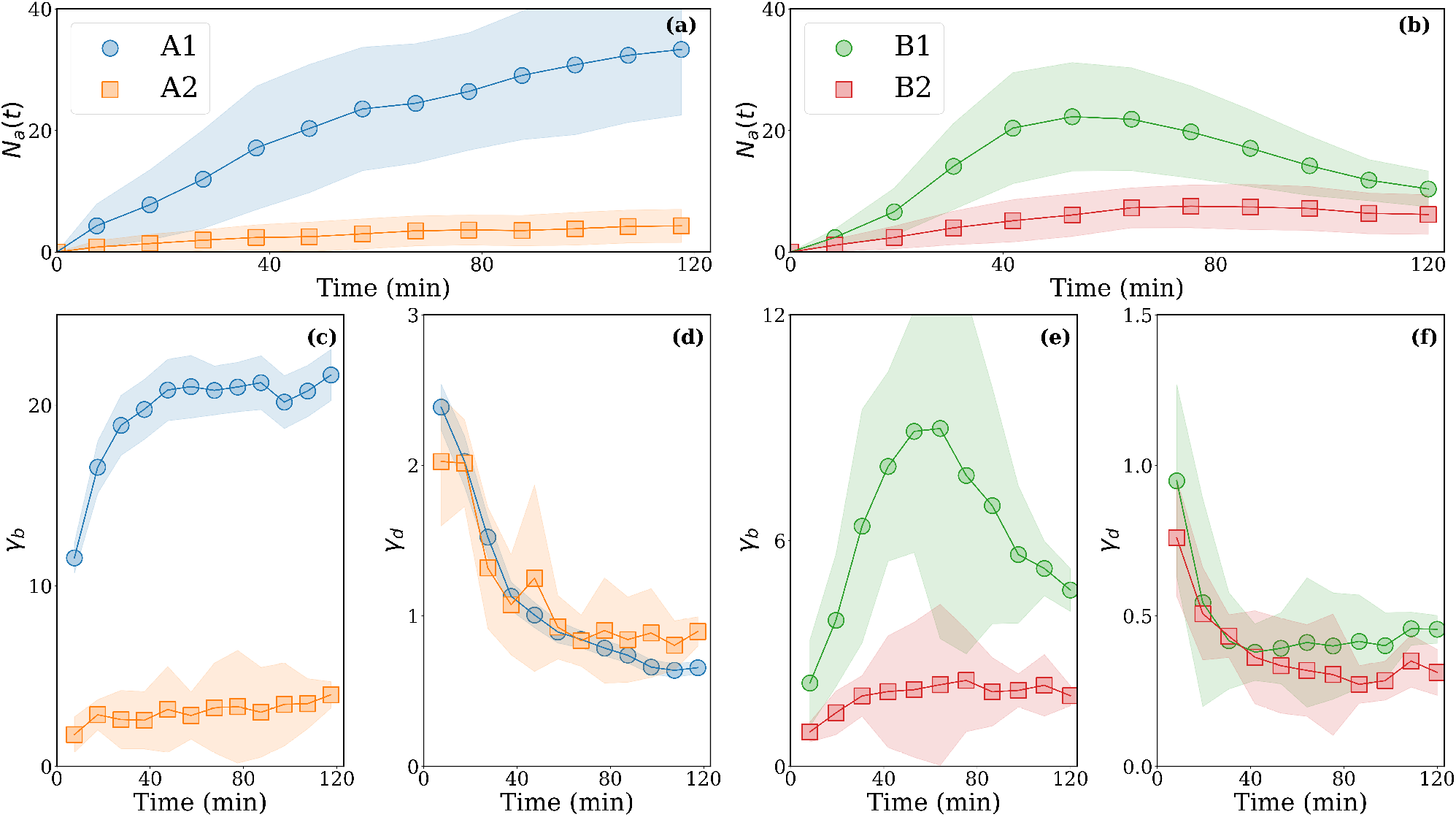
Average number of ants *N*_*a*_(*t*) and rates *γ*_*b*_ and *γ*_*d*_ obtained from the experimental datasets for the four colonies considered, two for experiment A (left) and two for experiment B (right). Patterns are shown for the full duration of experiments (120 minutes) to be compared with Fig. 2 showing the first 60 minutes of experiment. Error bars (shaded region) correspond to the standard deviation obtained by averaging over all the daily experiments carried out for each colony.

**FIG. 7.**
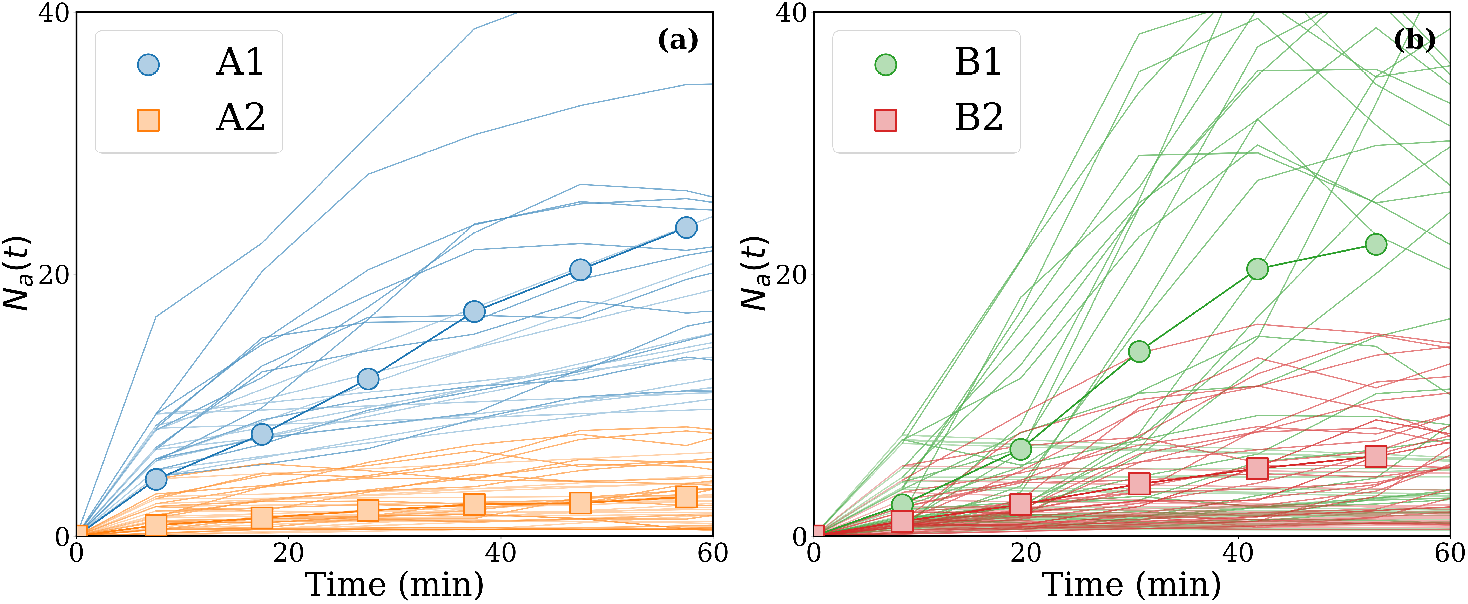
Average number of ants *N*_*a*_(*t*) obtained from the experimental datasets for the four colonies considered, two for experiment A (left) and two for experiment B (right). The light lines correspond to data of the daily experiments carried out for each colony.

Fig. 5 (left panel) shows how the optimal departure and return rates, and the corresponding optimal relative workforce derived from them, depend on colony size for a particular choice of *α*_*i*_ and *α*_*f*_ . As colonies grow larger, achieving 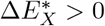 requires considerably larger departure rates. This leads to *γ*_*b*_ (blue circles) spanning a wide range of values (varying by nearly two orders of magnitude); instead, *γ*_*d*_ (red diamonds) remains comparatively stable, typically changing by less than a factor of two. This confirms again that the workforce load required to reach the maximum positive energetic balance increases with colony size. In practical terms, larger colonies need proportionally larger workforces (green triangles) to remain viable, and this increase in active foragers arises primarily from higher rates of recruitment from the nest, rather than from longer foraging trajectories.

In Fig. 5 (right panel), we compare again the two colony sizes, large (*N*_*col*_ = 500) and small (*N*_*col*_ = 100)), and examine how increasing the workforce load (*N*_*w*_*/N*_*col*_) influences the optimal departure and return rates. Here, vertical solid and dashed lines are included as a visual guide to mark the threshold where 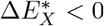 (solid red line), and the point where 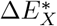 reaches its maximum value (dashed black line). Once again, these results show that engaging a larger workforce in the foraging process is achieved primarily by increasing *γ*_*b*_, that is, by launching new search trajectories at a higher rate. It is also noteworthy that *γ*_*d*_ consistently reaches a maximum and then declines once *N*_*w*_*/N*_*col*_ becomes sufficiently large (values above 0.45 for *N*_*col*_ = 100, and above 0.25 for *N*_*col*_ = 500). In these specific regions, the colony increases the number of active searchers by extending the duration of their search trajectories (decreasing the return rates) rather than by launching new ones. In small colonies (*N*_*col*_ = 100), however, this shift is much milder and occurs only at very high (likely unrealistic) values of *N*_*w*_. Within the more relevant region between the minimum (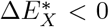 solid red line) and maximum (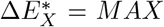 dashed black line) energetic balances, variation in (*γ*_*b*_)_*X*_ remains the dominant mechanism driving changes in workforce engagement, whereas adjustments in (*γ*_*d*_)_*X*_ play a comparatively minor role. This indicates that, for small colonies, modulation of the departure rate is the primary lever through which optimal foraging performance is achieved.

## IV. DISCUSSION

While there is substantial evidence that colonies of eusocial species can adjust their behaviour in response to environmental or other external perturbations, most documented cases focus on traits such as nest-site selection or resource exploitation. In contrast, the specific modulation of search and exploration patterns has received far less attention. In this regard, a central contribution of our work is to provide a predictive theoretical framework from which concrete, testable hypotheses emerge.

The metabolic rate of a colony is the aggregate of the metabolic rates of its individual members. When these individual rates are heterogeneous, one potential mechanism for regulating overall energy consumption is to adjust the relative proportions of high- and low-metabolism workers. Accordingly, colonies face nontrivial trade-offs in determining how much workforce they can allocate to foraging—the fundamental process through which they obtain energy from environmental resources. These trade-offs constrain both the potential rate of energy intake and the range of positive energetic balances that can be sustained as colonies increase in size. Together, these factors suggest that metabolic heterogeneity and limits on deployable foragers play central roles in shaping how colonies manage their energetic budgets during growth.

Our results reveal key behavioural adaptations that eusocial ant colonies may use to regulate their workforce *N*_*w*_ through the rates of worker departure (*γ*_*b*_) and return (*γ*_*d*_) from and to the nest. Within our modelling framework, we can hypothesize how a colony might adapt to an environment characterised by specific values of *α*_*i*_ and *α*_*f*_ . When these parameters make it relatively easy to maintain a positive energetic balance, colony fitness, and consequently colony size,should increase. However, as colonies grow, they face progressively tighter constraints on the feasible range of workers (i.e., *N*_*w*_ value) they can deploy. In other words, environments that allow colonies to thrive may simultaneously impose stricter limits on the workforce loads that can be sustained as colony size increases. Thus, while small colonies can remain viable by engaging only a modest number of active searchers, larger colonies increasingly require a greater proportion of active foraging workers in order to maintain a positive energetic balance. This scaling effect suggests that, for large colonies, maintaining a stationary reserve workforce becomes an advantageous, perhaps even necessary, evolutionary strategy.

In this context, our results provide further support for the idea that the presence of “lazy” or “inactive” workers [15–21], that is, individuals not heavily engaged in any specific task, should not be interpreted as inefficiency. Instead, they constitute a mechanism that maintains a permanent pool of workers ready to be mobilised when the colony’s energetic demands increase. This interpretation aligns with explanations for the sublinear scaling of metabolic rates observed in eusocial colonies and other social groups [19, 43–45]. According to this view, larger colonies favour a higher proportion of inactive workers, or equivalently, impose limits on the effective workforce load (from low-to high-metabolism individuals) as colony size increases. Empirical studies support this pattern: artificially increasing colony size leads to a higher proportion of inactive individuals [46], and in some species these can represent more than 50% of colony members [18, 23].

Ecological context further shapes how dynamic workforce modulation may benefit search efficiency in colonies. The spatial distribution of resources such as their average distance from the nest, degree of clustering, and the structural complexity of the habitat, is a major constraint on collective exploration and foraging efficiency [47]. Our results predict that environments where food detection is difficult (i.e., where energy intake *E*_*intake*_ or, equivalently *α*_1_, is reduced) should select for lower *γ*_*b*_ values and potentially higher *γ*_*d*_ values, resulting in a smaller available workforce *N*_*w*_. Colonies adapted to such low-resource habitats would therefore tend to remain small, relying on search strategies characterised by fewer, more asynchronous, and longer foraging trajectories. In contrast, habitats with higher resource availability should allow colonies to expand in size and adopt search strategies involving a larger number of synchronous foragers performing shorter trips. Under these conditions, increasing *γ*_*b*_ becomes the most efficient way to mobilise a larger workforce.

Most empirical studies examining the relationship between foraging activity and resource availability in eusocial species report patterns consistent with our predictions. Low resource density is typically associated with fewer but longer foraging trajectories, as well as a negative correlation between the distances travelled by searchers and the degree of group recruitment [11, 12, 48, 49]. Notably, studies such as [33, 50] have shown that these effects are largely independent of group size—provided it exceeds a critical threshold—supporting the idea that maintaining a reserve workforce can stabilize exploration strategies across a range of environmental conditions. In contrast, small groups of non-eusocial species (e.g., rodents) often exhibit the opposite pattern [51], highlighting the distinct collective dynamics that emerge in eusocial systems. Interpreting these results, however, is not always straightforward, as many reported foraging behaviours include phases of resource exploitation or are conducted under conditions where colonies may have been previously conditioned (explicitly or implicitly) by prior knowledge of resource locations.

If we assume that “collective cognitive programs” are themselves subject to group selection, a view that is increasingly supported in the ecological literature [15, 52–56], then our results offer a natural explanation for (i) the existence and functional relevance of inactive workers in large eusocial colonies, and (ii) the resulting metabolic scaling of colonies with their size. Together, these insights suggest that functional redundancy and the maintenance of workforce reserves are not incidental features of eusocial societies, but key components of their energetic stability, resilience, and evolutionary success.

More broadly, we argue that the redundancy principle [24] provides a particularly valuable lens for understanding collective behaviour in ecological systems. We hypothesize that this principle may operate across multiple tasks within the colony (not only in exploration), which would help explain the strong selection observed for the presence of task-unengaged individuals. Long-term monitoring of task allocation within colonies [57, 58], combined with controlled perturbations that disrupt one or several tasks and analysed within an energetic-cost framework such as the one developed here, would offer a powerful approach to testing this possibility.

## AKNOWDLEGMENTS

This research has been supported by the Spanish government through Grants No. PID2021-122893NB-C22 (DC and JC) and PID2021-122893NB-C21 (FB and PFL).

## Appendix A: Additional experimental results

In this Appendix we complement the empirical results discussed in Section III A to identify relevant questions related to their significance. In particular, we focus on how (i) the results obtained for the initial (search plus initial gathering) phase of foraging become modified as later phases are reached, and (ii) the implications and aspects related to the variability between colonies/experiments.

In Fig. 6 we show the values of the rates *γ*_*b*_ and *γ*_*d*_ obtained for the whole duration of the experiments (up to 120 minutes), so including later phases of the foraging process (essentially the end of the gathering phase and the pos-gathering period). It corresponds to a situation where the colony is actively exploiting (or has already exploited) food resources after the search phase, and so the colony behavior is a combination of (i) additional exploration for further resources, and (ii) relaxation dynamics once the probability to find them is low enough. Consequently, colonies exhibit in general a decreasing activity during that period (though the intensity of this strongly depends on the particular colony and the food conditions). This reflects into a certain decrease in the value of *γ*_*b*_ (so less active trajectories emerge out of the nest), but surprisingly *γ*_*d*_ does not increase significantly, or even go on decreasing during the final part of the experiments (except in the case of colony B1). Then, this stage is characterized by long trajectories of a few (specialized, probably) foragers trying to reach further regions of the arena, while the recruitment of new workforces remains at low levels.

Fig. 7, finally, shows the dispersion of results for *N*_*a*_(*t*) that we observe between individual experiments. While variability is considerable, it is remarkable that the characteristic regions occupied by each colony show low levels of overlap, which confirms our conclusion above that intercolony variability is much more significant than differences due to different food conditions and/or interday fluctuations. This means that adaptation responses of the groups are not immediate within the timescales of our experiments. It remains then to be explored experimentally how fast the colonies would be able to modify such strategies and/or what the main drivers are that would activate such adaptation.

## Appendix B: Effect of workforce modulation on search dynamics

As stated in Section II C, we computed random walk simulations of the search process with variable workforce to extract the values for the first-passage time *T*_*fp*_ (defined as the time required for any of the active foragers to find the resource) and the Collective Time Cost (*T*_*C*_, defined as the sum of the durations of all search trajectories in the simulation up to time *T*_*fp*_). These two are the only parameters in the energy balance from Eq. (5) that are computed numerically (the rest being chosen arbitrarily to facilitate visualization of our results).

By examining how ⟨*T*_*fp*_⟩ and ⟨*T*_*C*_⟩ depend on the rates *γ*_*b*_ and *γ*_*d*_, we gain additional insight into how search dynamics affect the colony’s energy balance. Figures 8 and 9 show the corresponding heatmaps for the same scenario illustrated in Fig. 3 of the main text, but now for ⟨*T*_*fp*_⟩ and ⟨*T*_*C*_⟩, respectively. These plots reveal that the first-passage-time *T*_*fp*_ (Fig. 8) is lowest when *γ*_*b*_ is small and *γ*_*d*_ is large. As *γ*_*d*_ increases, achieving comparable search efficiency requires increasingly larger *γ*_*b*_. Consequently, the balance between departure and return rates shapes the timescales over which food is found.

**FIG. 8.**
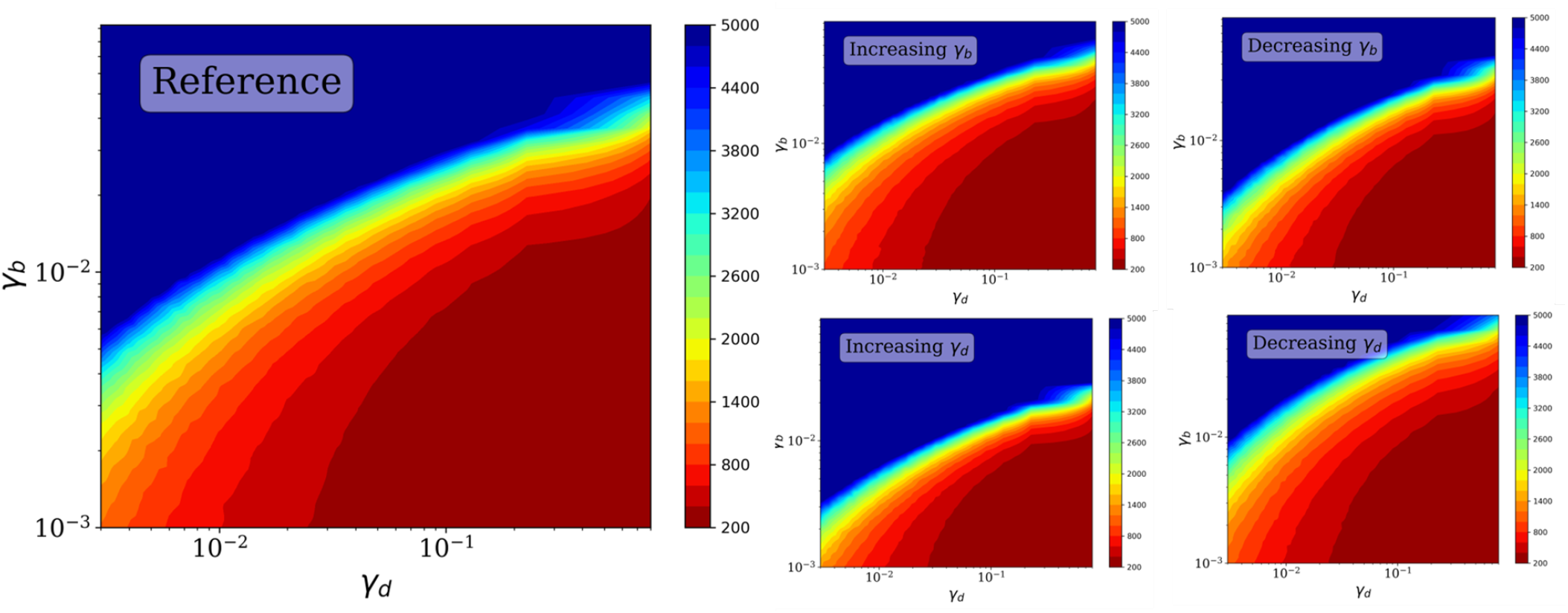
Heatmaps obtained from random-walk simulations for the the mean first-passage time, ⟨*T*_*fp*_⟩ through a target located at distance *d*_*r*_ = 10 (with *D* = 1 and *d*_*d*_ = 1) as a function of rates *γ*_*b*_, *γ*_*d*_. (Left): reference case where *γ*_*b*_, *γ*_*d*_ are kept constant during every simulation. (Right): Cases with either *γ*_*b*_(*t*) or *γ*_*d*_(*t*) increasing or decreasing by a factor of two during the search. Such an increase or decrease is linear with time and terminates after a characteristic time 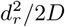, so after that time *γ*_*b*_ and *γ*_*d*_ will remain constant for the rest of the simulation.

**FIG. 9.**
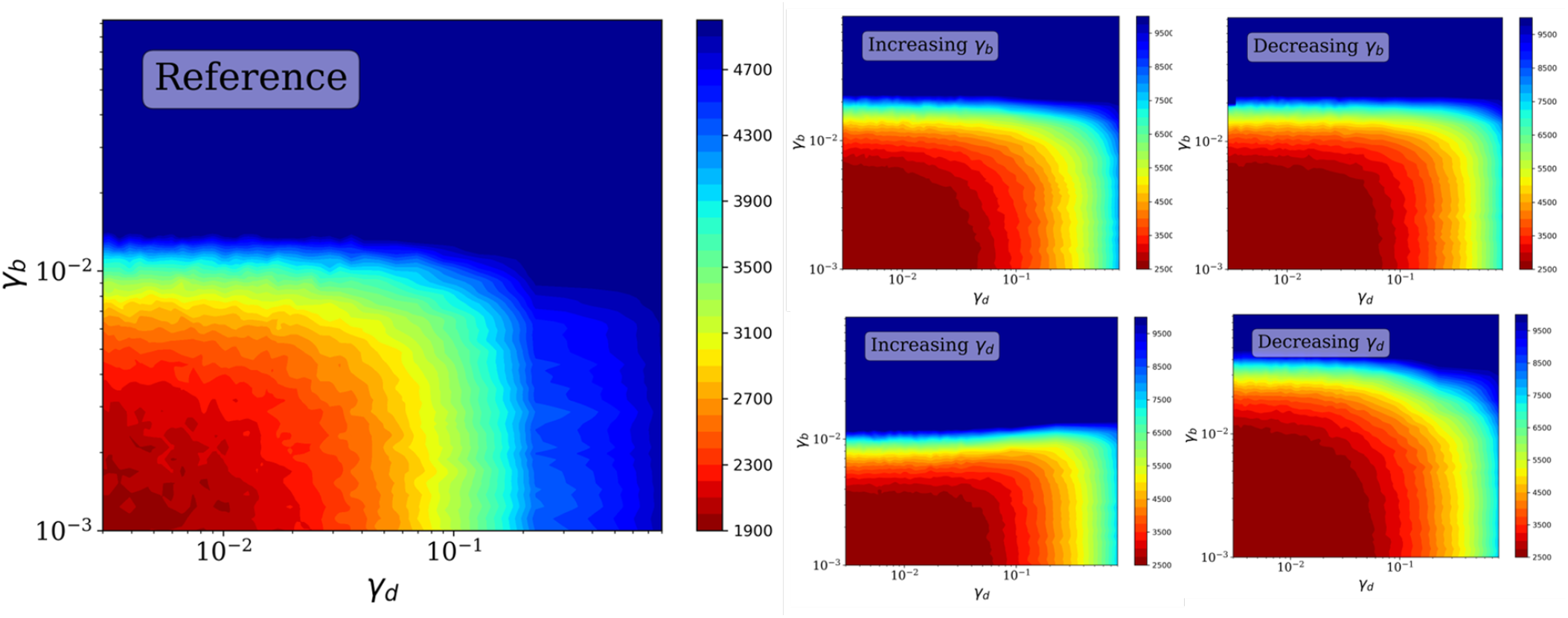
Heatmaps obtained from random-walk simulations for the collective search time, ⟨*T*_*C*_⟩, through a target located at distance *d*_*r*_ = 10 (with *D* = 1 and *d*_*d*_ = 1) as a function of rates *γ*_*b*_, *γ*_*d*_. (Left): reference case where *γ*_*b*_, *γ*_*d*_ are kept constant during every simulation. (Right): Cases with either *γ*_*b*_(*t*) or *γ*_*d*_(*t*) increasing or decreasing by a factor of two during the search. Such an increase or decrease is linear with time and terminates after a characteristic time 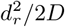, so after that time *γ*_*b*_ and *γ*_*d*_ will remain constant for the rest of the simulation.

In some regions of parameter space, the search effort becomes insufficient to locate the resource, leading to a pro-nounced increase in times required to find the resource (so, a decline in search efficiency). The regime characterized by low *γ*_*b*_ and *γ*_*d*_ reflects the idea that slowly increasing the number of scouts in the arena and subsequently keeping them synchronized there allows colonies to reduce resource detection times. Compared with stationary rates, non-stationary departure and arrival rates (right panels in Fig. 8) noticeably affect the first-passage-time efficiency. Increasing (decreasing) *γ*_*b*_ (*γ*_*d*_) over time slightly improves search times; however, the dominant effect is that increasing return rates (*γ*_*d*_) while decreasing departure rates (*γ*_*b*_) leads to an overall reduction in first-passage-time efficiency.

Opposite to the results for the first-passage times, the collective search time *T*_*C*_ is minimized (i.e., efficiency is maximized) in the regime of low (*γ*_*b*_) and low (*γ*_*d*_) (Fig. 9). Excessively large departure rates would impose an unnecessary workforce cost, whereas excessively large return rates would prevent collective search trajectories from remaining in the arena long enough to improve resource detection. Notably, decreasing *γ*_*b*_ and *γ*_*d*_ over time (right panels in Fig. 9) expands the region of low *T*_*C*_. In other words, the cost of collective search is reduced (i.e. search efficiency improves) when the system begins with a rapid turnover of trajectories that subsequently slows down, ensuring that workers in the nest are recruited only after previous trajectories have had sufficient time to travel far from the nest.

The energy balance described in the main text then reflects an interplay between these two components: the first-passage time, which favours resource-rich (or easy resource deteciton) and synchronous search, and the collective cost, which favours more resource-limited (or hard-to-find resources) asynchronous search strategies. This is further modulated by the basal energy consumption, which scales with colony size. Consequently, the energetic availability of the colony and the optimal energy gain Δ*E*_*X*_ studied throughout the article emerge from a nontrivial interaction among all these elements.

